# mTOR signaling regulates aberrant epithelial cell proliferative and migratory behaviors characteristic of airway mucous metaplasia in asthma

**DOI:** 10.1101/2024.02.12.579905

**Authors:** Katrina Kudrna, Elizabeth B. Staab, Evan Eilers, Paul Thomes, Shailendra Maurya, Steven L. Brody, Todd A. Wyatt, Kristina L. Bailey, John D. Dickinson

## Abstract

In asthma, the airway epithelium is hyperplastic, hypertrophied, and lined with numerous large MUC5AC-containing goblet cells (GC). Furthermore, the normal epithelial architecture is disorganized with numerous, what we here describe as, ectopic goblet cells (eGC) deep within the thickened epithelial layer disconnected from the lumenal surface. mTOR is a highly conserved pathway that regulates cell size and proliferation. We hypothesized that the balance between mTOR and autophagy signaling regulates key features of the asthma epithelial layer. Airway histological sections from subjects with asthma had increased frequency of eGC and increased levels of mTOR phosphorylation target-Ribosomal S6. Using human airway epithelial cells (hAECs) with IL-13 stimulation and timed withdrawal to stimulate resolution, we found that multiple key downstream phosphorylation targets downstream from the mTOR complex were increased during early IL-13-mediated mucous metaplasia, and then significantly declined during resolution. The IL-13-mediated changes in mTOR signaling were paralleled by morphologic changes with airway epithelial hypertrophy, hyperplasia, and frequency of eGC. We then examined the relationship between mTOR and autophagy using mice deficient in autophagy protein Atg16L1. Despite having increased cytoplasmic mucins, mouse AECs from Atg16L1 deficient mice had no significant difference in mTOR downstream signaling. mTOR inhibition with rapamycin led to a loss of IL-13-mediated epithelial hypertrophy, hyperplasia, ectopic GC distribution, and reduction in cytoplasmic MUC5AC levels. mTOR inhibition was also associated with a reduction in aberrant IL-13-mediated hAEC proliferation and migration. Our findings demonstrate that mTOR signaling is associated with mucous metaplasia and is crucial to the disorganized airway epithelial structure and function characteristic of muco-obstructive airway diseases such as asthma.

**Graphical Abstract Key Concepts:**

- The airway epithelium in asthma is disorganized and characterized by cellular proliferation, aberrant migration, and goblet cell mucous metaplasia.
- mTOR signaling is a dynamic process during IL-13-mediated mucous metaplasia, increasing with IL-13 stimulation and declining during resolution.
- mTOR signaling is strongly increased in the asthmatic airway epithelium.
- mTOR signaling is associated with the development of key features of the metaplastic airway epithelium including cell proliferation and ectopic distribution of goblet cells and aberrant cellular migration.
- Inhibition of mTOR leads to decreased epithelial hypertrophy, reduced ectopic goblet cells, and cellular migration.

## Introduction

Cells tightly regulate size, morphology, proliferation, and differentiation in response to nutrient availability and inflammatory or infectious cues. The barrier airway epithelium serves as a protective layer to the external environment. Consequently, airway epithelial cells must adapt to external signals for specialized functions. The identity and function of individual airway epithelial cells is now beginning to be fully characterized (1, 2). Yet during injury and repair, these cell identifications can be scrambled. The plasticity of the airway epithelium has been well described in response to injury and inflammation. In response to Type 2 airway inflammation, basal cells and secretory cells further differentiate into large mucous-secreting cells that are crammed with mucin granules, morphologically termed goblet cells. Epithelial metaplasia refers to an alteration of the cellular composition and cellular positioning in response to inflammation or repetitive injury and repair. In the cervical or gastric (3) epithelium, metaplasia has been heralded as a malignant precursor. However, in the airway epithelium, mucous metaplasia in diseases such as asthma and chronic obstructive pulmonary disease (COPD) leads to airway remodeling and airway lumenal occlusion by mucous plugs, culminating in airway obstruction. Metaplastic secretory cells are characterized by enlarged cytoplasm, ER stress, and protein and lipid synthesis (4–7). How airway epithelial cells regulate size, morphology, and movement in the context of dynamic exogenous signaling is not completely understood.

Mammalian target of rapamycin (mTOR) is a nutrient-sensing signaling protein that activates cell growth, metabolism, proliferation, and inhibits autophagy (8, 9). The mTOR pathway is a highly conserved kinase signaling system that sits at the hub of this network. Two distinct signaling complexes (mTORC1 and mTORC2) exist, with differing co-factors and phosphorylation targets. mTORC1 activation through phosphorylation of P70S6K and Ribosomal S6, and 4E-BP1 leads to increased protein synthesis and anabolic metabolism required for cell growth and proliferation (10–12). Inflammation, metabolic, and growth factor signaling cues converge upon mTORC1 activation, which inhibits autophagy through phosphorylation of ULK1 and ATG13 (13–16). Protein degradation is necessary for the cell to balance metabolic demands by recycling proteins to amino acids for new synthesis (17, 18). Autophagy is utilized in bulk protein breakdown (macro-autophagy or referred to as autophagy) in response to nutrient demands for new amino acids (19). We and others have reported that autophagy proteins are increased in the airway epithelium during mucous metaplasia *in vitro* (20, 21) and in diseased human airway samples (22). The common interpretation of these findings is that autophagy is activated during mucous metaplasia. Yet the abundant morphologic changes and protein synthesis that occur during mucous metaplasia suggests a more nuanced situation. Since mTOR is a powerful inhibitor of autophagy in most biologic systems, this suggests that earlier interpretations of autophagy protein abundance in relation to autophagy activity may be overly simplified. We hypothesized that the balance of mTOR and autophagy are important in the development and resolution of the disorganized metaplastic epithelial layer central to the airway remodeling found in human asthma.

## Methods

### Mice

Wildtype C57Bl6/j mice were purchased from Jax laboratories. Atg16L1 hypomorph mice (23, 24) were on the same background and were the gift of Dr. Thad Stappenbeck and Mouse Genetics Project, Wellcome Sanger Institute.

### Cells

Human AEC were derived from airways of donor lungs not suitable for transplant and seeded at 1.25 x10^5^ cells per 454.4mm^2^ surface area on 12 mm insert and differentiated on membrane supported inserts under air liquid interface (ALI) conditions as previously described (20, 21, 25). BEGM growth media (Lonza) was used for proliferation until cells were confluent, then media was transitioned to PneumoCult ALI media (Stem Cell). Mouse AECs were derived from wildtype and Atg16L1 hypomorph mice, expanded, and differentiated on membrane supported inserts under ALI conditions as previously described (23) with a seeding density of 8×10^4^ cell per surface area 133.73mm2 surface area.

#### Clinical samples

Histological sections were obtained from those who had a history of asthma and had died of fatal asthma attacks. Sections were provided courtesy of Dr. Derek Byers, Washington University. Non-asthma histological sections were obtained from those without a history of asthma whose lungs were rejected for transplant purposes. Sections were provided Dr. Kristina Bailey, University of Nebraska Medical Center. Characteristics are provided in **Table 1**.

**Table 1.**

| Table 1: |  |  |  |  |
| --- | --- | --- | --- | --- |
| Asthma sample | Age | Sex | Smoking status | Age of asthma diagnosis |
| IA1 | 67 | F | none | unknown |
| IA2 | 27 | M | no | age 24 |
| IA5 | 28 | M | no | unknown |
| IA6 | 42 | M | 1/2pk x 5yrs | age 41 |
| IA7 | 61 | F | none | age 57 |
| Non-asthma sample |  |  |  |  |
| MTS17 |  |  |  | NA |
| E17 | 35 | M | no | NA |
| B21 | 66 | M | no | NA |
| A16 | 76 | M | prior | NA |
| F18 | 68 | F | no | NA |

### Immunoblotting

mTOR proteins were sampled from human or mouse AECs. NP-40 buffer was used for lysis to preserve phosphorylation sites. The following antibodies were used for detection of protein on PVDF membranes following transfer: mTOR (Cell signaling), P70S6K total and T389 phosphorylation (Cell signaling), Ribosomal S6 total and S240, 244 phosphorylation (Cell signaling), ULK1 total and S757 phosphorylation (Cell signaling), and Stat6 total and phosphorylated (Cell signaling). Signal was quantified with Infrared-(IR) labeled secondary antibodies (LiCor) and normalized to total protein.

#### Immunohistochemistry

Slides made from formalin-fixed and paraffin-embedded tissue. They were incubated in xylene followed by Isopropanol. Slides were then washed in dH2O, followed by antigen retrieval with Trilogy (922P-06). Slides were then rinsed in TBST, followed by endogenous peroxidases quenching with 3% H2O2. Slides were then rinsed in TBST and blocked with 10% normal goat serum in PBS for 30 minutes at 37C in a humidity chamber. Slides were then incubated with primary antibody diluted in 0.5% BSA in PBS overnight at 4°C. Slides were then rinsed in TBST, followed by biotinylated goat secondary antibody in the same blocking buffer. After washing the slides in PBS, ABC reagent (Vector) was used, followed by chromagen substrate staining (DAB) (Vector). Slides were washed in PBS and counterstained with hematoxylin, washed in water and dehydrated with increasing concentration of ethanol incubations and then xylene.

### Immunofluorescence staining

Human and mouse AECs on membrane supports were harvested at the indicated time points. Basilar media was removed, and 200 microliters of 37-40°C 1% agarose was added to the apical surface. After cooling, the inserts were fixed in 10% formalin and embedded in paraffin. AECs inserts and human airway sections were processed for immunostaining using the same method at previously described (23). Antibodies for immunostaining included: mouse anti-MUC5AC 45M1 (ThermoFischer), rabbit anti Muc5ac (gift from Dr. Chris Evans), mouse anti Muc5b (gift from Dr. Burton Dickey), rabbit anti-Actin (Cell signaling), rabbit anti E-Cadheren (Cell signaling), rabbit anti-Cleaved Caspace 3 (Cell signaling), rabbit anti-p63 (Fitzgerald), rabbit anti-BrdU (Invitrogen). Image analysis was formed using set a common threshold image with ImageJ as previously described (21, 23, 25). Ectopic goblet cells (eGC) were defined as MUC5AC^+^ cells from human airway or hAEC sections that were not localized along the apical surface of the airway lumen. A blinded observer made the determination of eGC and normal GCs. Values were normalized by total GC and airway segment length. Normal GC were classified as being in direct contact with the lumenal surface, while eGC were classified if they had no direct contact with the lumenal surface.

### Cell migration assay

HAECs were expanded from passage 0 using BGEM media. At passage 1, 75,000 cells were seeded on a 24-well plate. When confluent, the 5% media was added to BGEM media ± IL-13 (10ng/mL) and rapamycin 250nM. Standard circular wounds were made in the center of cell layer. Wound closure was assessed by light field microscopy at 6, 24, and 30 hours timepoints or until complete closure. Wound area at each time point was normalized to the wound area at time 0.

### BrdU lineage tracing of human AEC

HAECs were seeded on membrane support inserts as previously described. When fully differentiated at approximately 21 days post ALI, hAECs were treated with ± IL-13 (10ng/mL) and rapamycin (250nM). All hAECs were treated with BrdU at final concentration of (10μM). A repeat BrdU treatment was then done at 48 hours. BrdU labeled media was removed at 96-hour post treatment. hAECs were harvested at day 8 post BrdU for immunostaining.

### Statistics

Statistical Analysis was performed using Prism 10. Methods for each analysis are described in the figure legend.

## Results

Metaplastic asthmatic airway epithelium is characterized by cellular hypertrophy, increased MUC5AC a fundamental disorganization of epithelial structure highlighted by ectopic goblet cells.

The airway in severe asthma has been previously characterized by airway remodeling with increased airway wall thickness, mucous metaplasia, cellular hyperplasia, and airway obstruction from mucus plugging (26–30). We examined airway sections from non-asthma controls and asthmatics to identify key features of the airway epithelial structure and localization with immunostaining of goblet cells with MUC5AC and cell borders with E-cadherin. Airway sections from non-asthma controls revealed the expected pseudostratified epithelial layer with intermittent MUC5AC+ goblet cells adjacent to the lumenal surface (**Fig. 1A panel #1**). In contrast, large airway sections from severe asthma revealed increased epithelial thickness, increased MUC5AC+ goblet cells (**Fig. 1A panels #1 and 2, and Fig. 1B, C)**, and occasional lumenal occlusion by a mucous plug in the small airways (<2mm in diameter) (**Fig. 1A panel #3**). Furthermore, the epithelial layer was fundamentally disorganized characterized by multiple layers of MUC5AC+ goblet cells. Many of the goblet cells were localized deep within the epithelial layer and without an outlet to the lumenal surface and were found in both large and small airway sections. These morphologically abnormally localized goblet cells were termed ectopic goblet cells (eGC). When quantified by fraction of eGC relative to all GC or number of eGC per airway segment, there was a significant increase in eGCs from those with asthma (**Fig. 1D, E**). However, eGCs were not unique to asthmatic airway sections. Some non-asthma control airway sections also contained scattered eGCs, though at a lower frequency. In both controls and severe asthmatic airway sections, the fraction of eGCs increased in a linear fashion with epithelial thickness of the corresponding airway segment (**Fig. 1F**).

**Figure 1:**
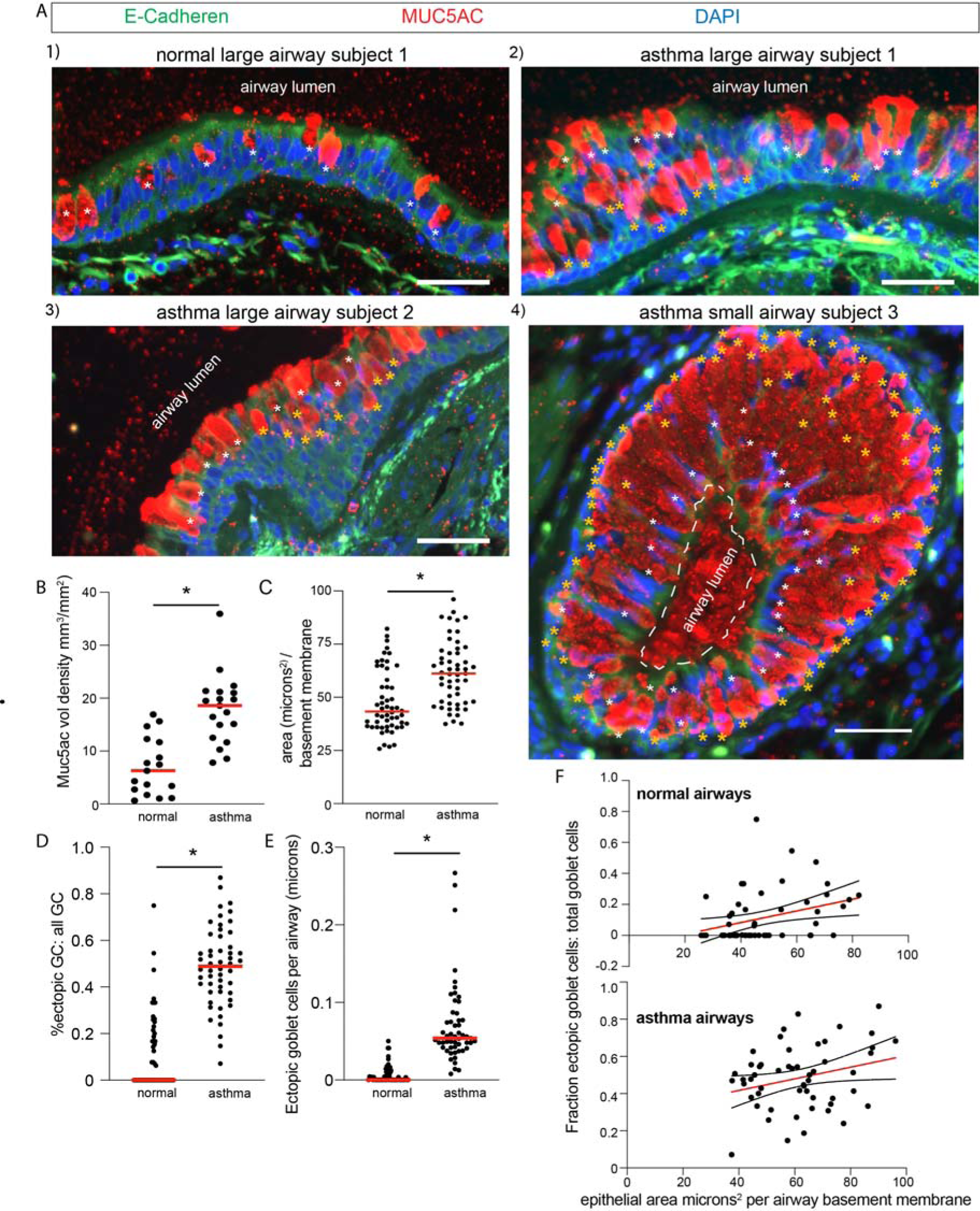
Airway epithelium from severe asthmatics is characterized by mucous metaplasia with disorganized goblet cell distribution. **A**) Representative airway immunostaining for MUC5AC and E-Cadherin: 1) Large airway section from non-asthmatic control, 2) Large airway section from #1 asthmatic sample, 3) Large airway segment from #2 asthma sample, 4) Small airway segment from #3 asthmatic sample. White asterisk marks airway luminal surface goblet cells, golden asterisk marks airway ectopic goblet cells (eGC). DAPI for nuclear counter stain. Scale bar equals 50 microns. **B**) Quantification of MUC5AC volume density per airway. N= 4 asthmatic airway and 5 normal airway donors. **C**) The total epithelial area micron^2^ per airway basement membrane segment was determined for non-asthma controls and asthmatic airways. N= 5 asthmatic airway and 5 normal airway donors with n=8 to 15 distinct images per donor. **D**) The fraction of ectopic goblet cells (eGC) relative to total goblet cells (**D**) and total cells (**E**) was calculated with n= 4 asthmatic airway and 5 normal airway donors with n=8 to 15 distinct images per donor airway. Unpaired T-test for statistical difference with * p<0.05. **F**) Simple linear regression comparison of airway epithelial area and fraction eGC relative to total GC. Upper panel is non-asthma controls and lower panel is asthmatic airways. Mean slope and 95% confidence intervals are shown.

### mTOR activation is a key feature of airway mucous cell metaplasia

We (20, 21), and others (31, 32) previously have shown that autophagosome protein LC3 II levels are increased during the development of mucous metaplasia in hAECs treated with IL-13. However, metaplastic airway epithelium is characterized by many of the central features of mTOR activation: epithelial protein synthesis, hyperplasia, and hypertrophy (**Fig. 1 and Supplemental Figure 1**). We hypothesized that cell signaling through mTOR may be associated with development of mucous metaplasia. We examined the airway sections from non-asthma controls and asthma for mTOR activation using p-RibS6, which regulates key mTORC1 signaling pathways in proliferation and protein synthesis. There was a significant increase in p-RibS6 staining in asthma sections compared to controls (**Fig. 2A-C**). This indicates that mTOR signaling is activated in the airway epithelium of asthma. We next sought to model the epithelial changes observed in airway sections from asthma. Airway epithelial cells (hAECs) were cultured on supported membranes and differentiated under air liquid interface (ALI) conditions. To model human asthma epithelial changes, hAECs were treated with human recombinant IL-13 for 7 days, then the cytokine was withdrawn to model resolution (**Fig. 3A**). IL-13 is known to induce cellular hyperplasia (33) and mucous metaplastic changes observed in human asthma (6, 7, 34–37). HAECs treated with IL-13 showed profound increases in cellular hypertrophy, epithelial area per airway segment, ectopic goblet cell fraction, and MUC5AC staining (**Supplemental Figure 1A-E**). These changes subsequently improved during resolution 7 days after withdrawal of IL-13.

**Figure 2:**
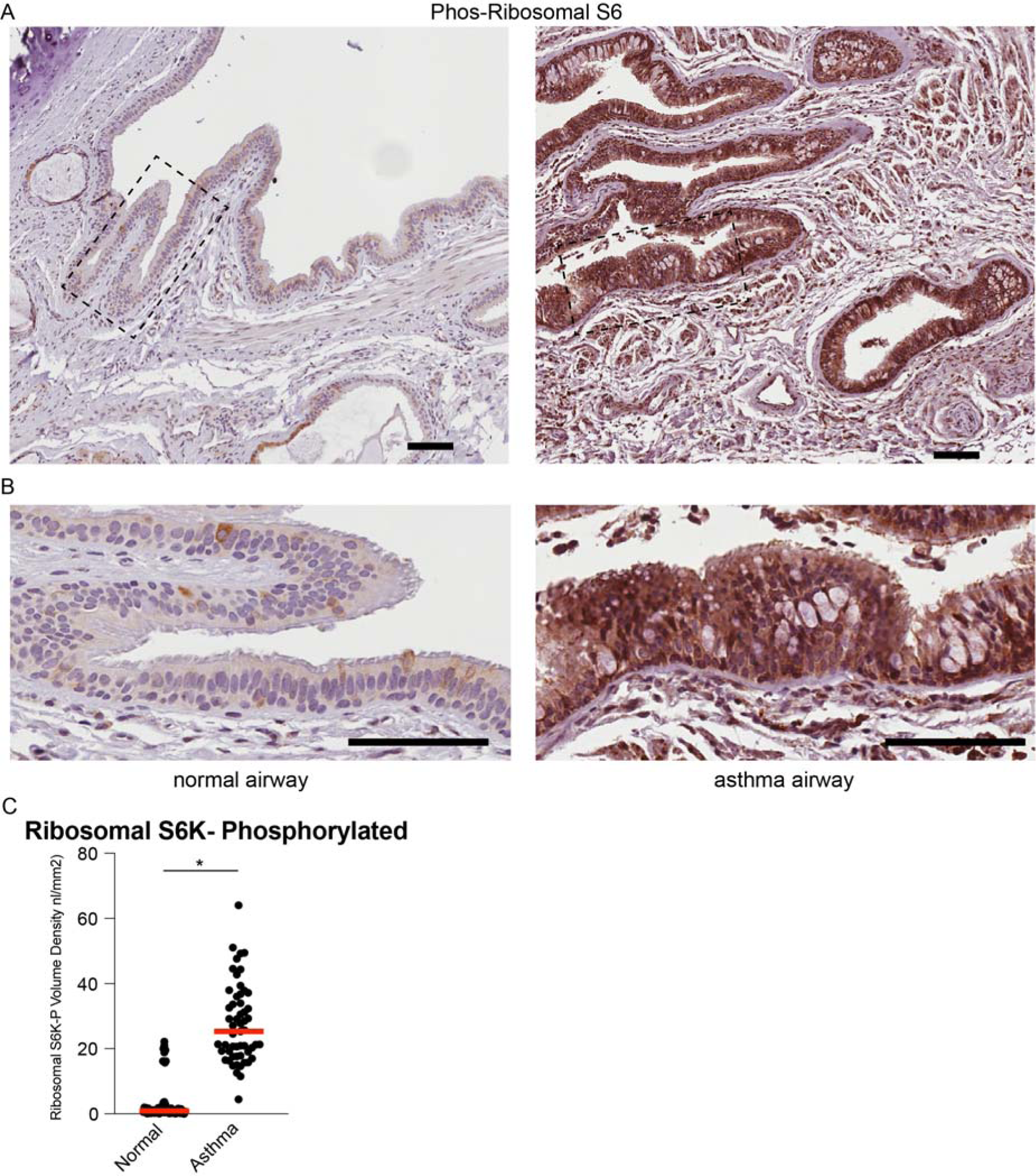
Increased mTOR signaling in asthmatic airway epithelium. **A**) Representative immunohistochemistry image of phosphorylated Ribosomal S6 (T240-244) from non-asthma controls and asthmatic airway sections. Scale bars equal 100 microns. Dashed box indicates area of higher magnification in **B**). Quantification of p-Ribosomal S6 levels normalized to volume density per airway (**C**). N=5 non-asthma controls and 5 asthmatic airway donors. 10-15 images per donor. Unpaired T-test for statistical difference with **\*** p<0.05.

**Figure 3:**
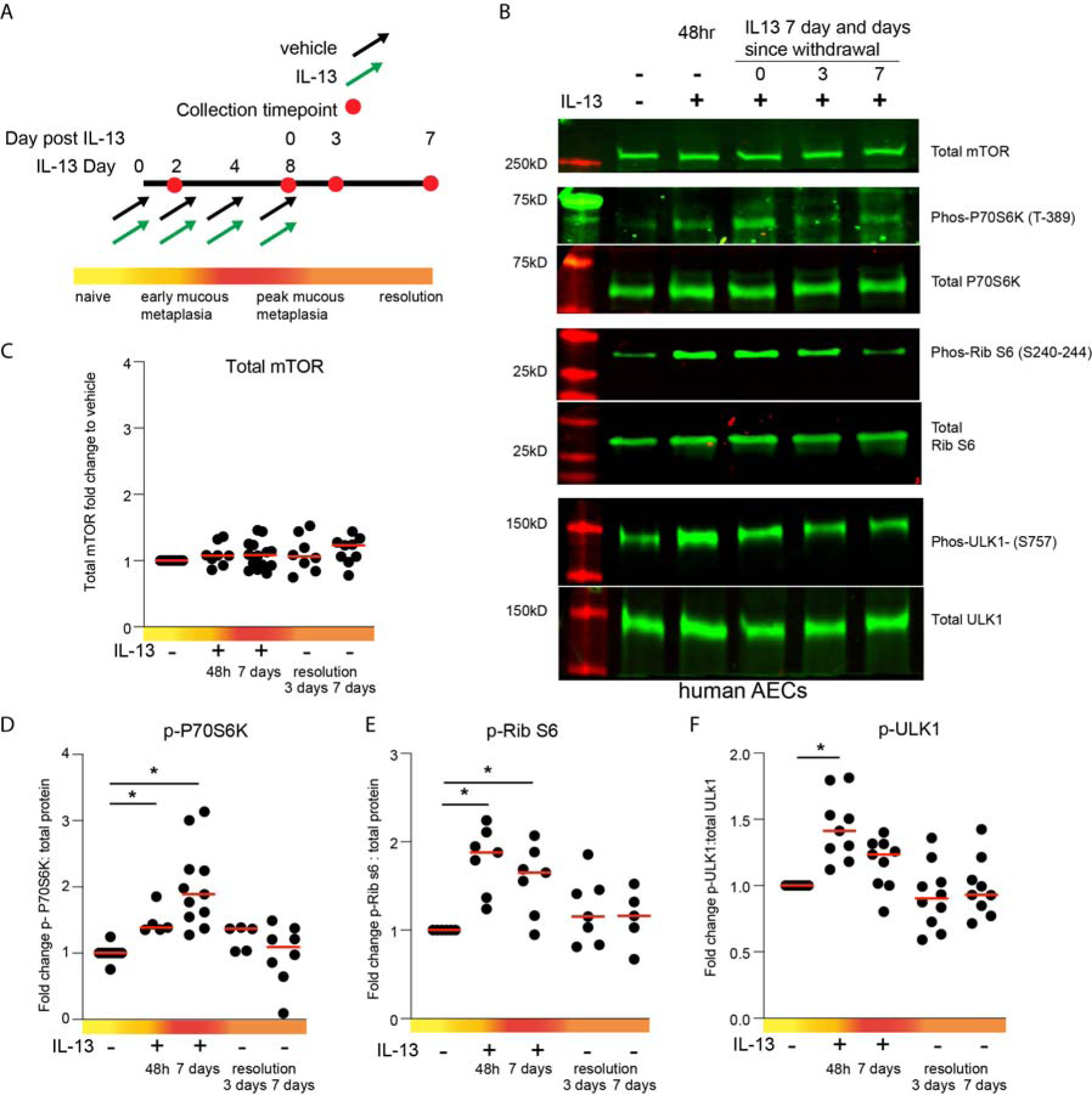
mTOR activity peaks early in IL-13-mediated mucous metaplasia, then diminishes during resolution. **A**) Scheme for IL-13-mediated mucous metaplasia in human airway epithelial cells (hAECs) and resolution according to timepoints after withdrawal of IL-13. **B**) Representative immunoblots for mTOR levels and total and phosphorylated Ribosomal S6 (T240-244), P70S6K (T389), and ULK1 (S757) during IL-13-mediated mucous metaplasia and resolution. **C-F)** Corresponding quantification of phosphorylated protein levels normalized to total protein lysate reported as fold change to untreated vehicle control (**B,C,D**). Phosphorylated proteins were normalized to non-phosphorylated protein. N= 5-8 per group from 3 unique donors. Kruskal Wallis for overall statistical difference with pairwise comparisons to vehicle control by Dunn’s multiple comparison test with significant difference. * p<0.05.

To characterize the dynamics of mTOR signaling, we evaluated multiple downstream mTOR signaling targets in (hAECs) under airway liquid interface (ALI) conditions during development and resolution of IL-13-mediated mucous metaplasia (**Fig. 2B).** We found that mTOR protein levels did not change during IL-13-mediated mucous metaplasia development or resolution (**Fig. 2 B, E**). However, downstream substrates of mTORC1 activation including phosphorylation of P70S6K, ribosomal s6 (RibS6), and ULK1 (**Fig. 2 B, D-F**) were significantly increased during early (48hr) and peak metaplasia timepoints (7 day of IL-13) and then decreased at day 7 of resolution. This suggests that mTOR signaling is activated early during IL-13-mediated mucous metaplasia, then significantly decreases during resolution. Assembly of ULK1 with ATG13 is required for the initiation of autophagy, and mTOR negatively regulates ULK1 by phosphorylation of reside S757 (15, 16). Consistent with increased mTOR signaling during early IL-13-mediated mucus metaplasia, we found an increase in phosphorylation of ULK1 at serine 757 in hAECs during early IL-13-mediated mucous metaplasia and a decrease in ULK-1 phosphorylation during resolution (**Fig. 2 B, F**). This indicates that autophagy is inhibited during early mucous metaplasia. However, during resolution, conditions are permissive for autophagy activation during low mTOR signaling.

### mTOR activation is upstream of autophagy in airway epithelial cells

We previously have shown that hAECs and mice deficient in autophagy have increased cytoplasmic MUC5AC levels (20, 23). To determine the relationship between mTOR signaling and autophagy during development and airway mucous metaplasia, we used mouse airway epithelial cells (mAECs) derived from Atg16L1 sufficient wildtype (WT) or Atg16L1 deficient mouse tracheas (Atg16L1^HM/HM^) cultured under ALI conditions with mouse recombinant IL-13 for 14 days (**Fig. 4A**). Similar to our *in vivo* findings (23), we found that mAECs from Atg16L1^HM^ mice had increased stored cytoplasmic Muc5ac and Muc5b levels (**Fig. 4 B-D**). We found similar IL-13-induced phosphorylation changes of P70S6K and ULK1 (**Fig. 4 E-G**). Interestingly, there was a trend toward lower p-Ribosomal S6 levels from Atg16L1^HM/HM^ AECs (**Figure 4 H, I**). mTOR directly phosphorylates ULK1, which inhibits autophagy (14), and also phosphorylates P70S6K, which then phosphorylates RibS6 to regulate protein synthesis and control of cell size (12). These data suggest that direct mTOR signaling remains intact when autophagy is blocked, and mucin granules accumulate. However, there may be a negative feedback loop that reduces ribosomal protein synthesis when autophagy is inhibited.

**Figure 4:**
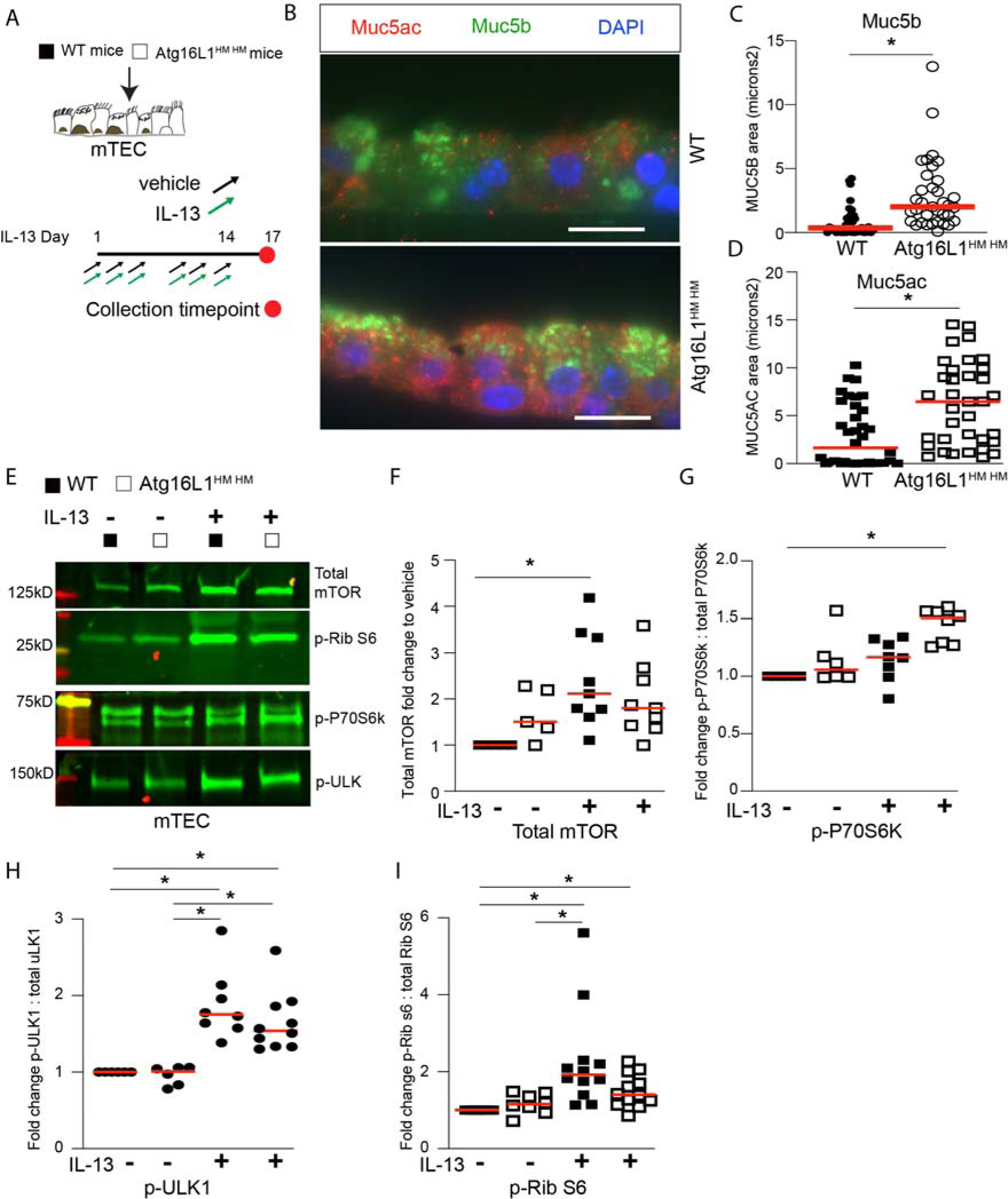
mTOR signaling independent of autophagy in IL-13-mediated mucous metaplasia. **A**) Mouse airway epithelial cells (mAECs) from Atg16L1 ++ (WT) or Atg16L1 deficient mice (Atg16l1^HM^ ^HM^) were expanded on inserts and differentiated under air liquid interface conditions. Then treated with mouse IL-13 for 14 days prior to collection. **B**) Representative immunostaining of Muc5ac and Muc5b in mouse AECs under ALI condition treated with IL-13 (10ng/ml) with corresponding quantification (**C,D**). Scale bar = 20 microns. Student T-test for significant difference <0.05. N= 4 inserts per condition from 2 independent tracheal preps and 8-12 microscopic images per insert. **E**) Representative immunoblots for total mTOR, p-ULK1 (S757), p-P70S6K (T389), and p-Rib S6 (T240-244) from mouse AECs. Corresponding quantification normalized to total protein reported as fold change to untreated vehicle (**F,G,H**). N=6-10 inserts per condition from 2 separate tracheal preps. For parts F-I, Kruskal Wallis for overall statistical difference with pairwise comparisons to vehicle control by Dunn’s multiple comparison test for significant difference. * p<0.05.

### mTOR activation is required for airway epithelial hypertrophy, hyperplasia, and ectopic goblet cell localization

To determine the role of mTOR in IL-13-mediated mucous metaplasia, we treated hAECs with vehicle control, IL-13, or IL-13 plus mTOR inhibitor, rapamycin, for 7 days (**Fig. 5A**). Rapamycin led to a significant decrease in p-RibS6 levels and autophagy protein SQSTM1 levels, indicating that mTOR was inhibited and autophagy was activated (**Fig. 5C, D**). We then examined hAEC morphology under these conditions by staining with MUC5AC for goblet cells and ACTIN to mark cell borders. We found mTOR inhibition led to a reduction in IL-13-mediated cellular hypertrophy, hyperplasia (**Fig. 5E, F, G, H**). There was no corresponding increase in cleaved caspase-3 due to mTOR inhibition (**Fig. 5I, J**) suggesting that cellular apoptosis could not explain the differences in cell size and number. Furthermore, the effect of mTOR inhibition on epithelial hypertrophy and hyperplasia was not due to reduction in basal cell populations. In fact, we found that p63^+^ basal cells were increased under IL-13 plus mTOR inhibition conditions (**Supplemental Figure 2 A, B**). We next examined the distribution of GC throughout the z axis of the airway epithelium as described in asthma airway sections in Figure 1. The fraction of eGC relative to total GC and overall MUC5AC staining was significantly reduced with mTOR inhibition (**Fig. 6A, B-D**). However, there was no change in Stat-6 phosphorylation between IL-13 and IL-13+rapamycin treated hAECs (**Supplemental Figure 2C-E**). This suggests that the IL-4/IL-13 receptor signaling is not directly dependent on mTOR. Overall, our findings suggest that mTOR signaling is central to the organization of goblet cells within the epithelial layer in response to IL-13 stimulation.

**Figure 5:**
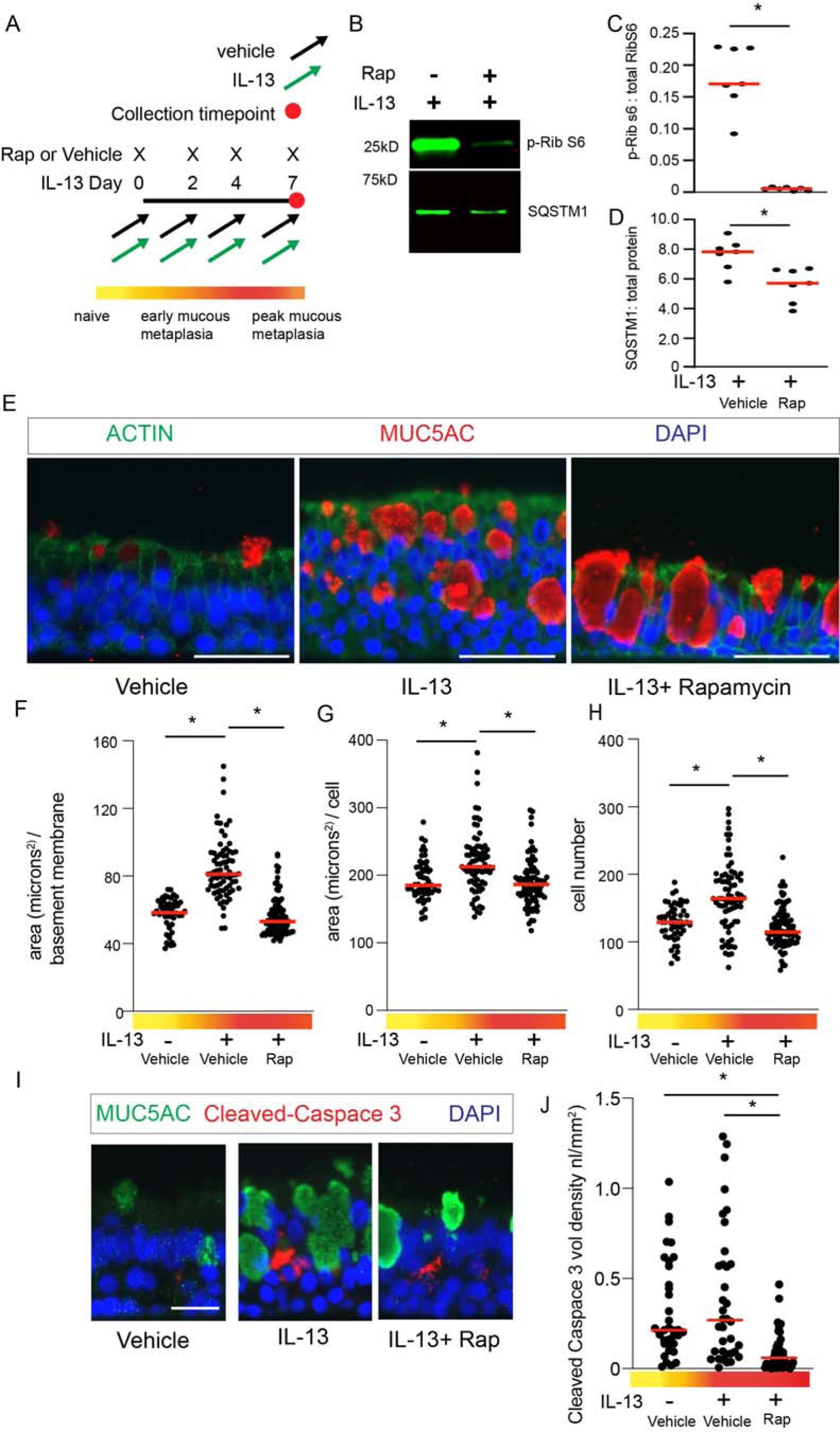
mTOR regulates airway epithelial hypertrophy and hyperplasia independent of apoptosis. **A**) Scheme for IL-13-mediated mucous metaplasia ± concurrent rapamycin treatment in differentiated hAECs under ALI condition. **B**) Representative immunoblots for phosphorylated RibS6 (T-240-244) levels and autophagy protein SQSTM1 with corresponding quantification normalized to total protein levels. Quantification of P-RibS6 normalized to total RibS6 (**C**) and SQSTM1 normalized to total protein (**D**). N=7 inserts from 3 independent donors. Student T-test for significant difference <0.05. **E**) Representative images of MUC5AC immunostaining for goblet cells and ACTIN immunostaining to outline cell borders of hAECs in untreated, IL-13 for 7 days, and IL-13+ Rapamycin**. F-H**) Quantification of epithelial area normalized to airway basement membrane, area per cell, and total number of cells. 10-12 microscopic fields from 6-8 inserts per condition from n= 3 unique normal airway donors. **I**) Representative images of MUC5AC and Cleaved Caspase-3 immunostaining. **J**) Quantification of Cleaved Caspace-3 immunostaining normalized to airway basement membrane. DAPI for nuclear counter stain. Scale bars =25 microns. 10-12 microscopic fields per donor from n= 4-5 inserts from each condition from 2 unique normal airway donors. ANOVA with Tukey multiple comparison test for statistical difference. **\*** p<0.05.

**Figure 6:**
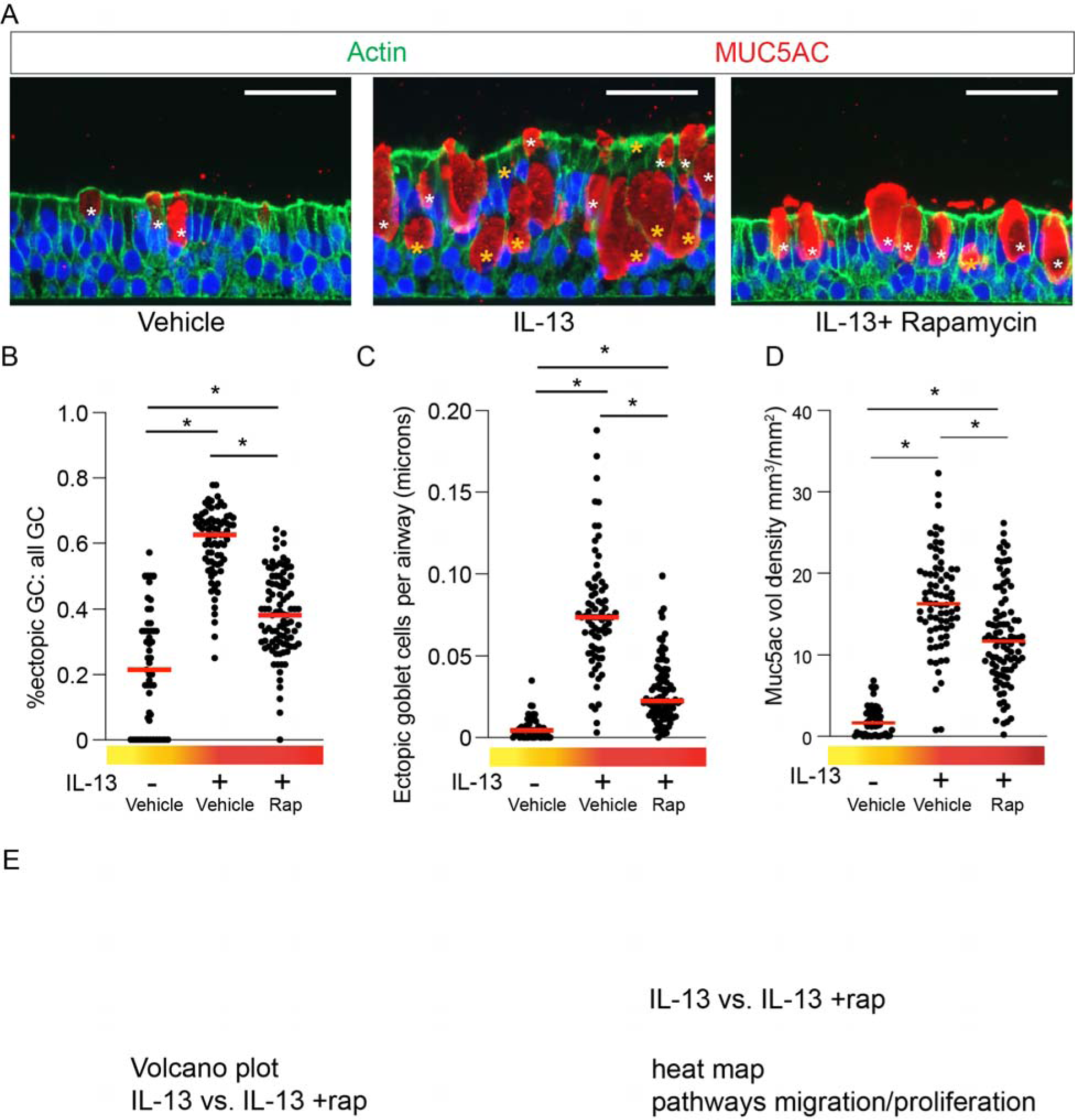
mTOR regulates ectopic goblet cells localization during IL-13 mediated mucous metaplasia. **A**) Representative images for MUC5AC and ACTIN imunostainingfrom vehicle, IL-13 7days, and IL-13 + Rapamycin 7 days hAECs. Surface GC were distinguished by a white * and ectopic GC were marked by a gold *. Scale bar =25 microns. Quantification of ectopic goblet cell (eGC) fraction per total goblet cells (**B**) and eGC per airway basement membrane length (**C**). Quantification of MUC5AC volume density immunostaining normalized to airway basement membrane length (**D**). Images from 10-12 microscopic fields from 7 inserts per condition from n= 3 unique normal airway donors. ANOVA with Tukey multiple comparison test for statistical difference. **\*** p<0.05.

### Aberrant epithelial proliferation and migration is dependent on mTOR

To examine the role of mTOR in epithelial proliferation and migration, we utilized hAECs to assess epithelial migration in response to wound assays. Short-term IL-13 stimulation led to a modest increase in cell migration. However, mTOR inhibition with rapamycin significantly decreased wound closure time in the presence of IL-13 (**Fig. 7A, B**). To further explore the impact of mTOR on proliferation and migration in fully differentiated hAECs, we utilized BrdU for lineage tracing of hAECs treated with vehicle control, IL-13, or IL-13+rapamycin for 7 days (**Fig. 7D**). Basal cells typically line the basement membrane and under control conditions, were frequently co-stained with BrdU. However, in response to IL-13, we found that BrdU-labeled cells were increased and more dispersed vertically in the epithelial layer away from the basement membrane. In contrast, mTOR inhibition with rapamycin reduced the total number of BrdU-labeled cells, and BrdU + cells were primarily concentrated along the basement membrane (**Fig. 7C, E, F**). This indicates that mTOR regulates epithelial cell migration and proliferation during IL-13-mediated mucous metaplasia and may contribute to the disorganized eGC distribution observed during airway remodeling in asthma.

**Figure 7:**
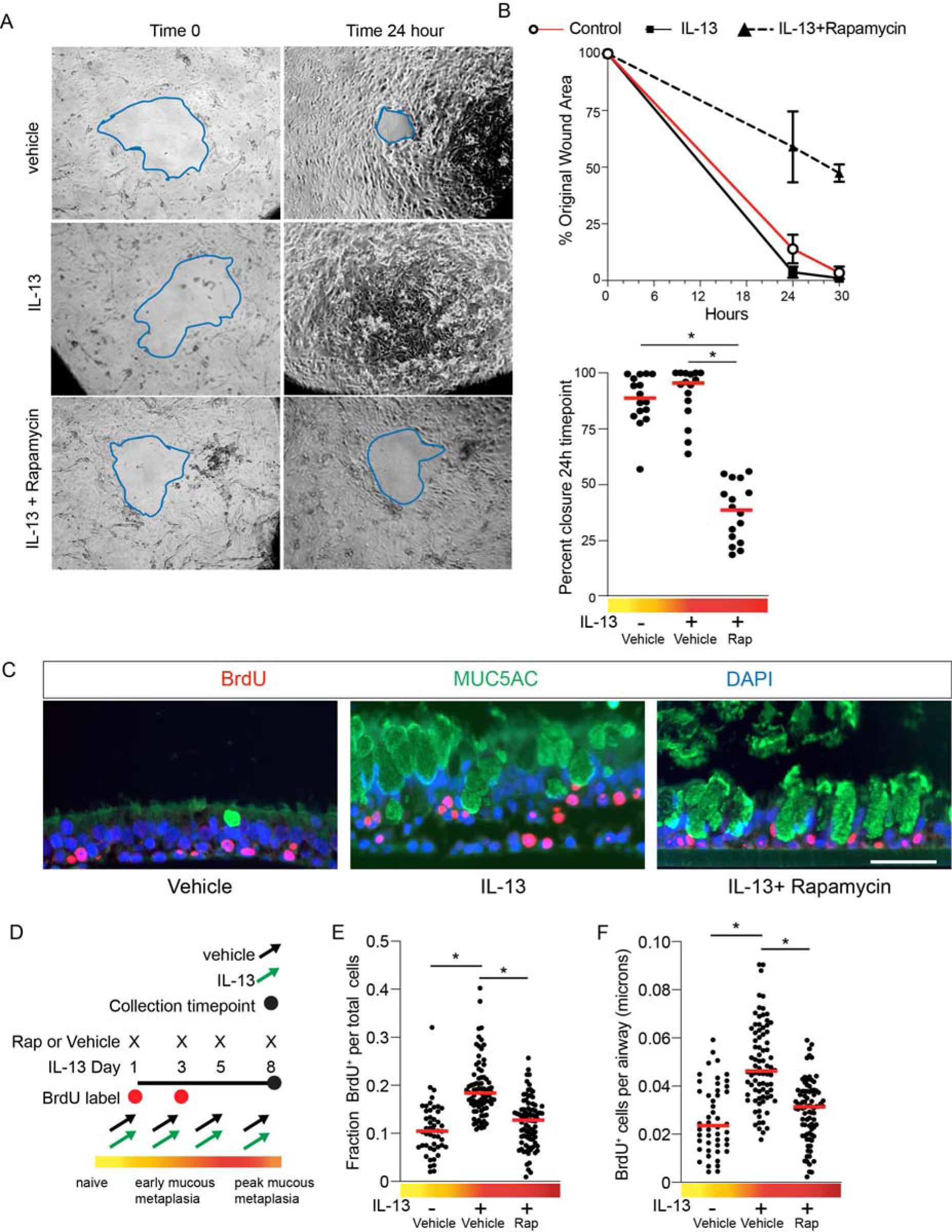
mTOR regulates two key features of airway mucous metaplasia; proliferation and migration. **A**) Representative light field microscopic images of hAEC at time 0 and 24 hours post circular wounding. Blue was used to pseudo-color the border of the wound area. **B**) Upper panel shows representative rate of wound closure as measured by percent closure compared to baseline wound area from one of four unique human donors. Lower panel depicts percent closure area at the 24 hour timepoint compared to time 0. N= 4 replicates from 4 unique hAEC donors. **C**) Representative MUC5AC and BrdU labeled hAEC cells for 3 conditions; Vehicle control, IL-13 7 days and IL-13 plus rapamycin 7 days. Scale bar =25 microns. **D**) Schematic showing timing of BrdU labeling during IL-13 mediated mucous metaplasia. **E**) Quantification of fraction of BrdU labeled cells compared to total cell number and **F**) quantification of number of BrdU normalized to airway segment baseline membrane length. N= 4 replicates from 3 unique hAEC donors with 10-15 microscopic images per airway segment. ANOVA with Tukey multiple comparison test for statistical difference. **\*** p<0.05.

## Discussion

Under homeostatic conditions, airway goblet cells release mucin granules into the airway lumen in a basal and stimulated fashion (38, 39), which contributes to mucociliary clearance of the airway. However, in muco-obstructive lung disease such as asthma, there is an increased number and size of goblet cells. Here we focused on airway epithelial hyperplasia, mucous metaplasia, and aberrant localization of goblet cells (ectopic goblet cells), which are a distinguishing feature of diseased airways including asthma. Our data suggest an mTOR-dependent, morphologically distinct population of goblet cells under metaplastic conditions that are aberrantly localized within a thickened epithelial layer and away from the lumenal surface. Recent work with single cell transcriptomics has led to the identification and characterization of subpopulations of airway epithelial cells with distinct roles, including distinct populations of goblet cells. During recurrent airway inflammation as in asthma, the normal differentiation paradigm is perturbed, leading to an altered differentiation program of distinct goblet cell populations (2). Histologically, this can be seen with epithelial hyperplasia, cellular hypertrophy, and mucous cell metaplasia (2, 27, 30, 40). Recently, the transcriptional changes by Type 2 stimulation (2), and specifically IL-13 (7), have been extensively characterized. Vast differentiation programing changes of secretory cell clusters derive metaplastic goblet cells from homeostatic goblet cells following IL-13 stimulation. These data point to broad epithelial-type 2 inflammatory cell interactions and paracrine growth factor signaling between epithelial cells. Our data shows that IL-13 activates mTOR signaling in the airway epithelium. Interestingly, mTOR inhibition had no impact IL-13 receptor signaling through stat6. This suggests a potential paracrine feed-back loop amongst epithelial cells, which directs the downstream mTOR signaling.

IL-13 stimulation recapitulates many of the morphologic features of the diseased epithelium in severe asthma(7). The surprising finding was that inhibition of mTOR reversed many of the morphologic changes: cellular hypertrophy, hyperplasia, and ectopic goblet cell distribution. We found that ribosomal S6, a key phosphorylation target of mTORC1 that regulates cell growth and protein synthesis, was significantly increased in the epithelium from severe asthmatics. Correspondingly, in human AECs we found that dynamic mTOR signaling increases during early IL-13 stimulation and drops precipitously during resolution after IL-13 withdrawal. mTORC1 signaling is known to play a key role in proliferation, cell growth, and protein synthesis in response to metabolic and inflammatory cues (10, 41). We found minimal caspase-3 staining following IL-13 stimulation and/or mTOR inhibition, suggesting that the loss of cell hypertrophy/hyperplasia with mTOR inhibition was not due to apoptosis. We detected small but significant increase in IL-13 dependent BrdU-labeled hAECs that was mTOR dependent. This suggests that IL-13 acts as an epithelial cell growth factor for proliferation and confirms the findings of previous studies (33, 42).

Our morphological findings from asthma airway sections reveal that ectopic goblet cells (eGC) are proportional to the airway epithelial thickness. We hypothesize that the aberrant localization of these eGC was due to increased migration in an mTOR-dependent fashion. Previously it has been shown that IL-13 induces increased cellular migration (43) in wound assays for undifferentiated hAECs. Our data found an increase in epithelial wound closure at 24hr in the IL-13 treatment group. More importantly, mTOR inhibition with rapamycin lead to significant decrease in migration of IL-13 treated undifferentiated hAECs. The significance of mTOR in hAEC migration was confirmed in differentiated AECs using BrdU labeling. Not only did rapamycin-treated, IL-13 stimulated hAECs have fewer BrdU-labeled cells, indicating a loss of proliferation, but rapamycin treatment led to a loss of the apparent lumenal mobility of BrdU-labeled cells. While our focus has been on the role of mTORC1 in directing features of airway epithelial remodeling during asthma, recent data suggest that mTORC2 may play a role in regulating cellular orientation and migration (44–46). Future studies are warranted to dissect the unique roles of mTORC1 and mTORC2 in regulating airway epithelial proliferation and migration in normal development and disease.

The first report on the role of mTOR in asthma mouse models was 20 years ago. It was observed that systemic administration of mTOR inhibitor SAR943 significantly reduced OVA-mediated airway inflammation, epithelial proliferation, and number of goblet cells (47). Other studies have confirmed the importance of mTOR in Type 2 models of asthma in mice (48, 49). Interestingly, in our findings in human asthma airway sections, IL-13-stimulated differentiated hAECs and mAECs all consistently demonstrate that mTOR-mediated phosphorylation of RibS6 is significantly increased. In contrast, a recent study (50) reports the oppositive finding with decreased p-RibS6 in asthma airway sections, mouse lungs, and undifferentiated hAECs. However, there are several distinctions worth noting between these two studies. First, in that study, hAECs were undifferentiated at time of IL-13 stimulation. Second, the authors used an antibody to a different phosphorylation site pRibS6 (T236), which may account for the divergent findings. Indeed, earlier reports suggested that RibS6 could be phosphorylated at T235 and T236 independent of mTOR-mediated P70S6K signaling (51). Furthermore, our study confirmed mTOR activation with multiple different downstream phosphorylation targets, including P70S6K, RibS6, and ULK1. Finally, IL-13 and other asthma-related growth factors are mitogenic with associated increases in epithelial cellular hypertrophy, hyperplasia, and goblet cell hyperplasia. These processes all rely on profound transcriptional and protein synthesis, which exemplify mTOR activation in all cell types.

Our study has several limitations worth noting. While the IL-13 model for type 2 asthma is reproducible, it does not capture the complex inflammatory milieu typical of asthmatic airway, nor do our findings address non-atopic asthma. Secondly, we rely on pharmacologic loss of function studies with rapamycin. Genetic gain/loss of function studies may be able to confirm our findings and tease out specific mTOR substrates that are critical in the airway epithelium.

## Conclusion

Asthma is a complex human disease that involves recurrent and eventually persistent airway obstruction due to the impact of inflammation and infection on airway epithelial cells, stromal cells, and local inflammatory cells. Here we describe a key significant role of mTOR in the airway epithelium and suggest that mTOR signaling plays a vital role in the disorganized epithelial morphology found in asthma airways.

## Acknowledgements

JDD is supported by NHLBI-R01HL157269.

TAW is supported by P50-AA030407, U54 OH010162, and I01 BX005886. TAW is the recipient of a Research Career Scientist Award (IK6 BX005962) from the Department of Veterans Affairs.

KB is supported by ACORN P50 AA030407 and VA-I01 BX006049

## Abbreviations

Airway epithelial cells (AECs), airway liquid interface (ALI), Bromodeoxyuridine (BrdU), Chronic Obstructive Pulmonary Disease (COPD), goblet cell (GC), interleukin 13 (IL-13), mammalian target of rapamycin (mTOR), Protein S6 Kinase B2 (P70S6K), Ribosomal protein S6 kinase B1 (Rib S6).

---

**Supplemental Figure 1:**
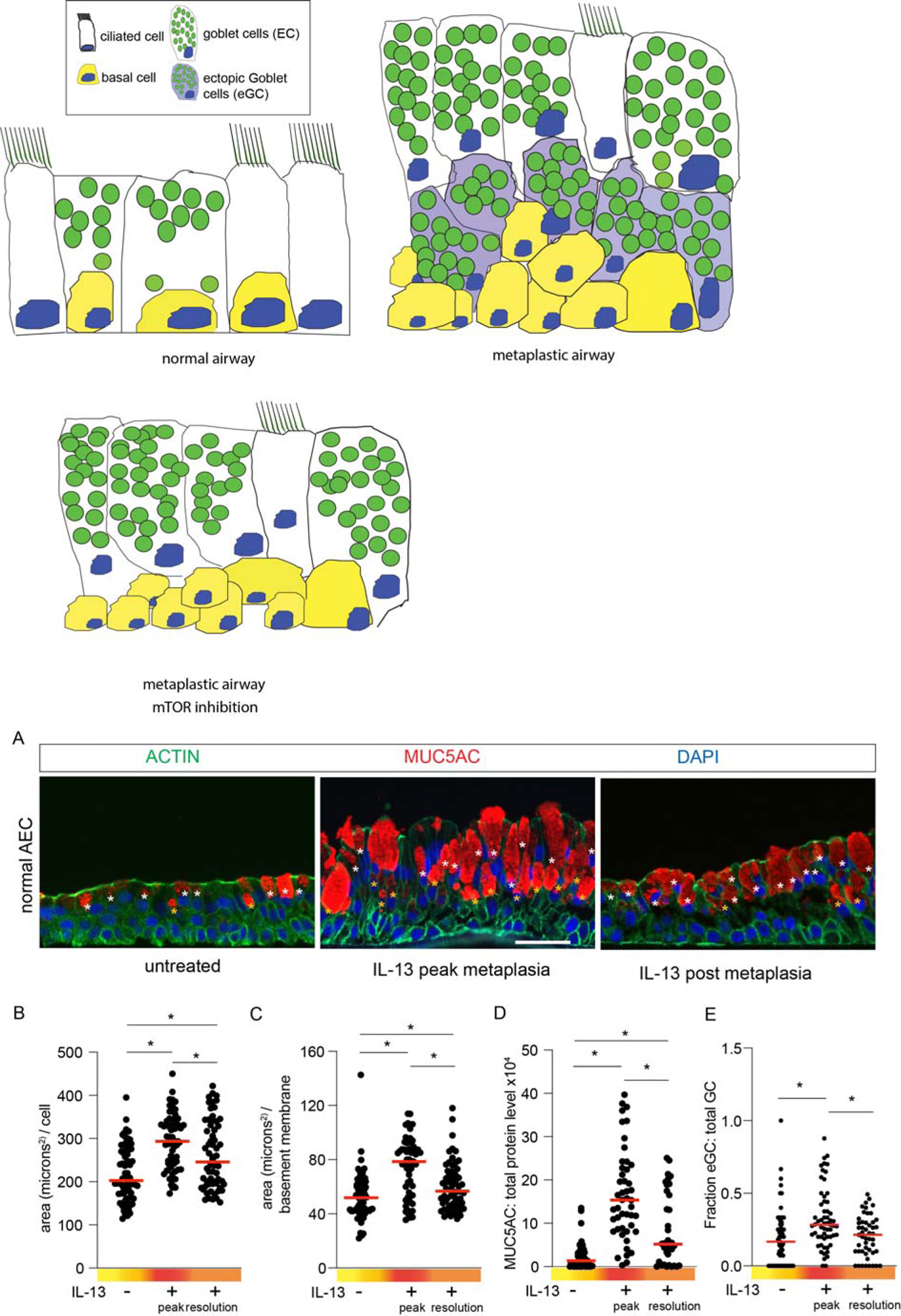
**A**) Representative images MUC5AC and ACTIN immunostaining of normal human airway epithelial (hAECs) in untreated, metaplastic (IL-13 10ng/mL for 7 days), and post-metaplastic (7days post withdrawal) conditions. DAPI for nuclear counter stain. Scale bars =50 microns. White asterisk marks airway luminal surface goblet cells, golden asterisk marks airway ectopic goblet cells (eGC). Quantification of area per cell (**B**), total cell area per airway segment (**C**) and MUC5AC volume density per airway segment (**D**). **E**) Quantification of fraction ectopic GC from total goblet cells. 10-20 microscopic fields per donor from n= 4 replicates from 4 unique normal airway donors. ANOVA with Tukey multiple comparison test for statistical difference. **\*** p<0.05.

**Supplemental Figure 2:**
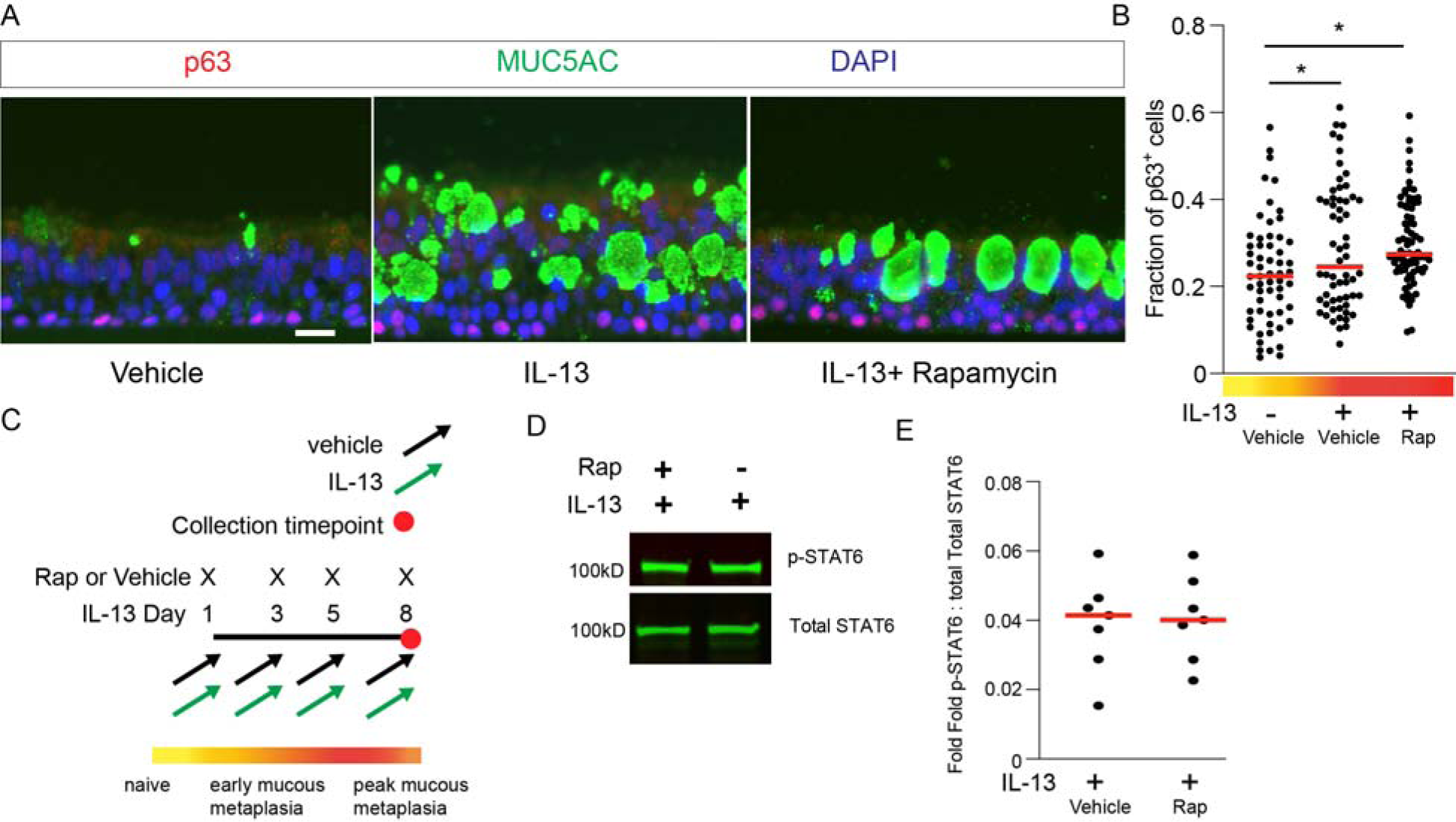
mTOR inhibition does not block IL-13-mediated Stat 6 phosphorylation or p63+ basal cell expansion. **A**) Representative immunostaining of basal cell marker, p63, and MUC5AC. DAPI for nuclear counter stain. Scale bar equals 20 microns. **B**) Quantification of fraction of p63+ cells from total DAPI stained cells per airway image. N= 9-10 microscopic images from 6-8 inserts per condition from 3 unique hAEC donors. **C)** Scheme for IL-13-mediated mucous metaplasia ± concurrent rapamycin treatment in differentiated hAECs under ALI condition. **D**) Representative immunoblots for Stat-6 and phosphorylated Stat6 (Tyr 641) with corresponding quantification (**E**) of P-Stat6 normalized non-phosphorylated Stat-6 and total protein levels. N=7 inserts from 3 unique hAEC donors. ANOVA with Tukey multiple comparison test for statistical difference. **\*** p<0.05.

